# Relationships between endogenous circadian period, physiological and cognitive parameters and sex in aged non-human primates

**DOI:** 10.1101/2021.05.24.445429

**Authors:** Clara Hozer, Fabien Pifferi

## Abstract

The biological clock exhibits circadian rhythms, with an endogenous period *tau* close to 24h. The circadian resonance theory proposes that lifespan is reduced when endogenous period goes far from 24h. It has been suggested that daily resetting of the circadian clock to the 24h external photoperiod might induce marginal costs that would accumulate over time and forward accelerate aging and affect fitness. In this study, we aimed to evaluate the link between the endogenous period and biomarkers of aging in order to investigate the mechanisms of the circadian resonance theory. We studied 39 middle-aged and aged *Microcebus murinus*, a nocturnal non-human primate whose endogenous period is about 23.1h, measuring the endogenous period of locomotor activity, as well as several physiological and behavioral parameters (rhythm fragmentation and amplitude, energetic expenditure, oxidative stress, insulin-like growth factor-1 (IGF-1) concentrations and cognitive performances) in both males and females. We found that aged males with *tau* far from 24h displayed increased oxidative stress. We also demonstrated a positive correlation between *tau* and IGF-1 concentrations, as well as learning performances, in males and females. Together these results suggest that a great deviation of *tau* from 24h leads to increased biomarkers of age-related impairments.

**Summary Statement:** This manuscript describes how the misalignment between the endogenous circadian period and the 24h of the environmental periodicity impacts aging biomarkers and accelerates cognitive decline.

## Introduction

Circadian rhythms are biological rhythms with an endogenous period *tau* close to 24h. They are ubiquitous in living organisms and are thought to be highly adaptive features (Sharma, 2003; Sheeba and Sharma, 1999). They help the organism synchronizing its functions with optimal phases of the daily fluctuating environment and are supported by the circadian clock, the timekeeping system which synchronizes every day activity with the periodic external light/dark cycles. The circadian clock controls a wide range of biological processes such as metabolic rate, sleep/wake cycles, hormone production, or temperature profile (Barclay et al., 2012; Kalsbeek et al., 2014; Karatsoreos et al., 2011; Kyriacou and Hastings, 2010; Reinke and Asher, 2016; Richards and Gumz, 2013; Scheer et al., 2013; Wright et al., 2012). The central pacemaker is located in the suprachiasmatic nuclei (SCN), in the hypothalamus (Moore, 2013). The circadian rhythms are generated by a set of clock genes (*Clock, Bmal, Cry* and *Per* among others) that interact with each other in transcriptional and posttranscriptional loops in order to maintain the rhythm’s periodicity (Duong et al., 2011; Gekakis et al., 1998; Lande-Diner et al., 2013; Yu et al., 2002). The adaptive advantage of the biological clock in physiological and behavioral outputs is well known, and demonstrated in numerous studies. For example, chronic jet-lag impairs memory and learning and leads to various disease states such as inflammatory disorders (Craig and McDonald, 2008; Gibson et al., 2010; Preuss et al., 2008). Analysis of clock genes mutations also provides valuable insights into the damaging impact of circadian disorganization: *Clock* mutation induces metabolic dysfunctions as well as sleep alterations (Naylor et al., 2000; Turek et al., 2005); the *tau* mutation in the circadian regulatory gene casein kinase-1∊ deeply affects cardiovascular and renal integrity (Martino et al., 2008).

Besides, the use of targeted disruptions or mutations of clock genes enables to highlight the implication of the circadian clock in aging. Deficiency for clock genes invariably leads to age-associated pathologies, and/or reduced lifespan (Zhu et al., 2009). For example, mice deficient for *Bmal1* gene exhibited sarcopenia, cataracts, or organ shrinkage, among others (Kondratov et al., 2006). Mice deficient for *Clock* gene developed age-specific cataracts and dermatisis, and displayed about 15% reduced lifespan compared to wild type (Dubrovsky et al., 2010). In addition, the restoration of proper circadian rhythms and extended lifespans in aged rodents transplanted with fetal SCN is a great proof of the major implication of the biological clock in aging dysfunctions (Cai et al., 1997; Li and Satinoff, 1998; Viswanathan and Davis, 1995). The complexity of the link between circadian clock and aging lies in the fact that aging and the biological clock influence each other. Indeed, aging also impacts circadian features by fragmenting the activity patterns, reducing rhythms’ amplitude, shifting the onsets, or altering sleep-wake cycles (Carrier et al., 2002; Farajnia et al., 2012; Nakamura et al., 2015; Yoon et al., 2003).

Aging may be involved in the relationship between lifespan and biological clock, described through the circadian resonance theory. This theory postulates that the deviation of *tau* from 24h is negatively correlated with longevity: the more *tau* gets far from 24h, the shorter the lifespan. Several studies provided evidence to support this assumption in fruit flies, mice and primates (Hozer et al., 2020; Libert et al., 2012; Pittendrigh and Minis, 1972; Wyse et al., 2010). Interestingly, the mechanisms underlying this postulate are up to now unknown. They could be related to daily costs of synchronization: in order to cope with the deviation of *tau* from 24h, the organism would engender daily metabolic costs to reset the clock. These costs, accumulated over the lifetime would forward accelerate aging processes and impact longevity. In a recent study, it has been shown that a mismatch between endogenous and external rhythms induced higher resting body temperature, as well as higher resting metabolism and lower cognitive performances in a small primate, suggesting a daily additional energetic expenditure when *tau* and the environment periodicity do not resonate (Hozer and Pifferi, 2020). However, to our knowledge, the consequences of these costs on aging patterns have not been yet investigated. A study reported that long-lived mice exhibited a 24h endogenous period at young and old age but did not investigate its influence on aging biomarkers (Gutman et al., 2011). In the present study, we investigated the transversal relationship between the endogenous period and aging parameters such as metabolic activity, aging cellular biomarkers (growth hormone IGF-1, oxidative stress) and cognitive performances in middle-aged and aged individuals.

We performed these experiment on a small Malagasy primate, the gray mouse lemur (*Microcebus murinus*), a strictly nocturnal species with a mean endogenous period lying around 23.1h (Hozer & al, 2020). Its metabolism is highly dependent on photoperiod, exhibiting high levels during the long-day photoperiod, whereas food scarcity during the short-day photoperiod compels low metabolism and sexual rest (Génin and Perret, 2003; Schmid and Speakman, 2000; Vuarin et al., 2014). These changes are only driven by the photoperiod, even in captivity (Languille et al., 2012; Perret and Aujard, 2001). The gray mouse lemur has gained interest these last decades, because of the age-related impairments it displays. Indeed, the gray mouse lemur exhibits age-related spontaneous neurodegenerative diseases and cognitive deficiencies such as Alzheimer’s disease or memory losses similar to those found in humans (Hozer et al., 2019; Languille et al., 2012).

## Material & Methods

### 1) Ethical statement and housing conditions

All mouse lemurs studied were born in the laboratory breeding colony of the CNRS/MNHN in Brunoy, France (UMR 7179 CNRS/MNHN; European Institutions Agreement no. E91–114.1). All experiments were performed in accordance with the Principles of Laboratory Animal Care (National Institutes of Health publication 86–23, revised 1985) and the European Communities Council Directive (86/ 609/EEC). The research was conducted under the approval of the Cuvier Ethical Committee (Committee number 68 of the ‘Comité National de Réflexion Ethique sur l’Expérimentation Animale’) under authorization number 12992-2018011613568518 v4.

All cages were equipped with wood branches for climbing activities as well as wooden sleeping boxes. The ambient temperature and the humidity of the rooms were maintained at 25 to 27°C and at 55% to 65%, respectively. When they are not involved in experimental protocols, animals in facilities are exposed to an artificial photoperiodic regimen consisting of alternating periods of 6 months of long day (Light-Dark (L:D) 14:10) and 6 months of short days (L:D 10:14), in order to ensure seasonal biological rhythms (Perret and Aujard, 2001). The experiment focused on animals tested during the two last months of the short-day season.

### 2) Experimental procedure

39 individuals (15 females and 24 males) were monitored. Mean ages were 6.23±0.74 and 6.45.3±1.12 years for females and males respectively. They followed the same experimental protocol during the last months of the short-day season: their locomotor endogenous period was measured in free-running conditions (*i.e.* total darkness) during 15 days. Blood and urine were sampled. The metabolic activity was measured using a calorimetry system during 5 days. Afterwards, the animals performed 2 cognitive tasks during the last 3 days of experiment: learning in a visual discrimination task and working memory in a spontaneous alternation task were evaluated. Animals were fed at random times of the day with fresh fruits and a homemade mixture, corresponding to an energy intake of 25.32 kcal.day^−1^ (see Dal-pan et al., 2011 for details). Body mass were monitored every 3 weeks and averaged over the whole short-day season.

### 3) Indirect calorimetry system

Metabolic activity was recorded using an indirect calorimetry system (Oxymax, Colombus Instruments Inc, Columbus, Ohio, USA). Animals were housed in individual metabolic cages during 5 days (with one day for acclimation before measurements). Oxygen consumption (VO_2_), carbon dioxide production (VCO_2_) and energy expenditure (*Heat* = 3.815 × *VO*_2_ + 1.232 × *VCO*_2_, see Lusk, 1924) were recorded continuously. VO_2_ and VCO_2_ were expressed as a function of the whole body mass (mL.h^−1^.kg^−1^) and Heat in kcal.h^−1^.

### 4) Locomotor activity monitoring

Recording of locomotor activity (LA) was obtained by infra-red sensors (for details, see Matikainen-Ankney et al., 2019). LA data were summed in intervals of 1 minute and expressed in arbitrary unit (a.u.). Activity onset was defined as the first six successive bins where activity was greater than the mean LA during the resting phase. Individual mean endogenous period was assessed derivating the slope of the line connecting onsets with Clocklab software (Actimetrics, Evanston, IL, USA) and expressed in hours. We only used the 7 last days, in order to avoid the transition phase between light-dark cycles and free-run. The rhythm amplitude of locomotor activity corresponded to the mean daily difference between maximum and minimum locomotor activity. Rhythm fragmentation (or IV = Intradaily variability) was calculated using following formula (Witting et al., 1990):

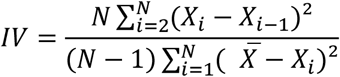

with *N* = total number of individuals, *X*_*i*_ = locomotor activity at time *i*.

### 5) Oxidative stress and insulin-like growth-factor 1 measurements

Mouse lemurs’ blood was collected 3h before light off, in order to minimize daily fluctuations between individuals. Two hundred μL of blood were taken via the saphenous vein, and collected in tubes containing EDTA. Blood samples were then centrifuged at 2000g at 4°C for 30 minutes and plasma was collected and stored à −80°C. We assayed plasma 8-hydroxy-2’-deoxyguanosine (8-OHdG) levels (OxiSelect™ Oxidative DNA Damage Elisa kit, Cell Biolabs Inc.) and insuline-like growth-factor 1 (IGF-1) concentrations (IGF-1 ELISA assay, Demeditec Diagnostics GmbH, Kiel, Germany) in ng.mL^−1^ in duplicate on two aliquots of 25 μL. For technical reasons (too low volume sample), 7 values could not be measured for 8-OHdG and 14 for IGF-1.

### 6) Visual discrimination task (learning)

The visual discrimination task was first described by Picq et al. (2015), inspired from apparatus designed by Lashley (1930) for rodents and based on visual discrimination. In the present study, it was conducted over a two-day period, 3h hours before the light extinction. The first day is dedicated to animals’ habituation to the set-up. The discrimination test takes place on the second day. The animal is introduced in a big squared vertical cage through an opening in the wall, to an elevated starting platform. It has to jump onto one of two landing platforms, fixed below on the opposite wall. A hole centered behind the two landing platforms leads to a nesting box behind the cage. On each landing platform, a visual *stimulus* helps discriminating the left and right platforms. One of these visual clues is the positive *stimulus*, the other is the negative one. At each trial, the landing platform with the positive clue is kept fixed and leads to the nesting box, whereas the platform with the negative clue is mobile and toggles when the animal jumps on it, such a way that it falls on a cushion pillow, to prevent any injury. At each trial, the location of the positive and negative stimuli on the right or left landing platform is randomized. The animal was given 30 trials to reach the success criterion consisting in 8 jumps on the positive platform within 10 consecutive trials. For each animal, we measured two parameters: -the number of trials needed to reach the success criterion, -the total success rate to the task (number of correct jumps/number of total jumps). We also split the individuals into two groups, those who reached the criterion and those who did not (yes/no) (for more details, see Picq et al., 2015). 9 individuals failed the habituation phase or refused to jump during the discrimination task and were thus excluded from the analysis.

### 7) Spontaneous alternation task (working memory)

The spontaneous alternation maze is a hippocampal dependent task, that testes spatial working memory. It is a useful test that reduces stress, in particular because it does not require food deprivation. The test was conducted 3h before light extinction. The maze is composed of 4 wood right-angled arms (50 cm long, 32 cm high and 16 cm wide), with different spatial cues on each arm. The mouse lemur is placed in the intersection of the 4 arms and is allowed to navigate freely in the maze during 20 minutes, using the external cues. Entries into every arm were noted (4 paws had to be inside the arm for a valid entry). We considered a total spontaneous alternation if an animal entered the four different arms consecutively. We considered a partial alternation if the animal entered three different arms over 4 consecutive entries. Percentages of total and partial spontaneous alternation were calculated according to following formula: (number of alternations)/(total number of arm entries−3)×100.

### 8) Statistical analyses

Several parameters are known to covary with body mass, as it influences body composition, particularly body fat (Engstrom et al., 2006; Onder et al., 2006); therefore, body mass was included in all statistical models. We conducted linear models between every parameter and the endogenous period. Sex, age, body mass and *tau* were integrated as fixed effects. Age was incorporated as a categorical variable, individuals being separated into three categories: those aged 5-6 years, those aged 6-7 years and those aged over 7 years. We also performed a generalized binomial model to investigate whether endogenous period could be related to the reach of the criterion (yes/no). We considered all interactions two by two between fixed parameters. We selected the best models using a backward procedure by calculating Akaike’s Information Criterion (AIC), conserving only models with the smallest AIC, but keeping the fixed effect *tau.* Residuals normality was verified using Shapiro-Wilk tests. All data that had not a normalized distribution were log-transformed.

## Results

The endogenous period was not correlated with age, but displayed significant divergence between sexes: females exhibited a shorter period than males (females: 22.73±0.35h, males: 23.22±0.32h, Table 1 and Fig. 1). Regarding oxidative stress, an interaction has been found between *tau* and sex, and between *tau,* sex and age (Table 1). Indeed, in males over 7 years of age, the 8OHdG plasma level was negatively correlated with *tau* (R^2^=0.62, Fig. 2C). However, in females, 8OHdG concentrations increased with *tau* (R^2^=0.26, Fig. 2B). *Tau* was positively related to IGF-1 (R^2^=0.22, Table 1 and Fig. 3), as well as the success rate at the learning task (Fig. 4A), regardless of the sex. This result is corroborated by the fact that individuals who did not reach the success criterion have significantly shorter *tau* (22.80±0.40h) compared to those who did reach the criterion (23.21±0.32h, Fig. 4B). However, no relationship was found between the number of errors before reaching the success criterion and the endogenous period (R^2^=0.22, Table 1). Finally, neither rhythm amplitude, nor rhythm fragmentation or mean locomotor activity were correlated to *tau* (Table 1).

**Table 1:** Effects of *tau,* age, sex and body mass on rhythmic, metabolic, cellular and cognitive outputs

|  | *Tau* |  | *Sex* |  | *Age* |  | *Tau\*Sex* |  | *Tau\*Age* |  | *Age\*Sex* |  | *Tau\*Sex\*Age* |  | *Body mass* |  |
| --- | --- | --- | --- | --- | --- | --- | --- | --- | --- | --- | --- | --- | --- | --- | --- | --- |
|  | *F* | *p* | *F* | *p* | *F* | *p* | *F* | *p* | *F* | *p* | *F* | *p* | *F* | *p* | *F* | *p* |
| **Tau** | - | - | 19.48 | **0.0001** | 0.54 | 0.58 | - | - | - | - | - | - | - | - | - | - |
| **Amplitude** | 0.59 | 0.44 | - | - | - | - | - | - | - | - | - | - | - | - | 3.66 | **0.04** |
| **Fragmentation** | 0.75 | 0.10 | - | - | - | - | - | - | - | - | - | - | - | - | 10.61 | **0.002** |
| **Activity** | 0.38 | 0.54 | - | - | - | - | - | - | - | - | - | - | - | - | 1.86 | **0.04** |
| **VO2** |  |  |  |  |  |  |  |  |  |  |  |  |  |  |  |  |
| Resting phase | 0.04 | 0.85 | 0.25 | 0.62 | 2.59 | 0.1 | 1.60 | 0.22 | 0.02 | 0.98 | 2.04 | 0.15 | - | - | - | - |
| Active phase | 0.37 | 0.55 | 1.02 | 0.32 | 0.52 | 0.60 | - | - | - | - | 5.73 | **0.01** | - | - | - | - |
| **Heat** |  |  |  |  |  |  |  |  |  |  |  |  |  |  |  |  |
| Resting phase | 0.02 | 0.89 | 0.34 | 0.57 | 0.14 | 0.87 | 2.86 | 0.11 | 3.15 | 0.07 | 2.02 | 0.16 | - | - | - | - |
| Active phase | 0.31 | 0.58 | 1.64 | 0.21 | 0.22 | 0.80 | - | - | - | - | 4.11 | **0.03** | - | - | - | - |
| **Log(8OHdG)** | 1.04 | 0.32 | 0.02 | 0.88 | 1.54 | 0.24 | 6.79 | **0.02** | 1.31 | 0.29 | 1.06 | 0.36 | 6.24 | **0.007** | - | - |
| **IGF-1** | 9.54 | **0.005** | 4.24 | 0.05 | - | - | - | - | - | - | - | - | - | - | - | - |
| **Total alternation** | 0.01 | 0.7 | 0.04 | 0.84 | 0.13 | 0.73 | - | - | - | - | - | - | - | - | - | - |
| **Partial alternation** | 0.13 | 0.72 | 0.18 | 0.69 | 0.67 | 0.44 | - | - | - | - | 1.38 | 0.28 | - | - | - | - |
| **Success rate** | 11.58 | **0.002** | 0.57 | 0.45 | 6.29 | **0.007** | - | - | - | - | 6.06 | **0.008** | - | - | - | - |
| **Errors to criterion** | 3.23 | 0.11 | 0.001 | 0.98 | 2.20 | 0.17 | - | - | - | - | 2.02 | 0.19 | - | - | - | - |
| | $\chi^2$ | *p* | $\chi^2$ | *p* | $\chi^2$ | *p* | $\chi^2$ | *p* | $\chi^2$ | *p* | $\chi^2$ | *p* | $\chi^2$ | *p* | | |
| **Reached the criterion (Y/N)** | 14.54 | **0.0001** | 6.76 | **0.009** | 3.65 | 0.06 | - | - | 8.84 | **0.003** | 2.47 | 0.12 | - | - | - | - |

**Figure 1:**
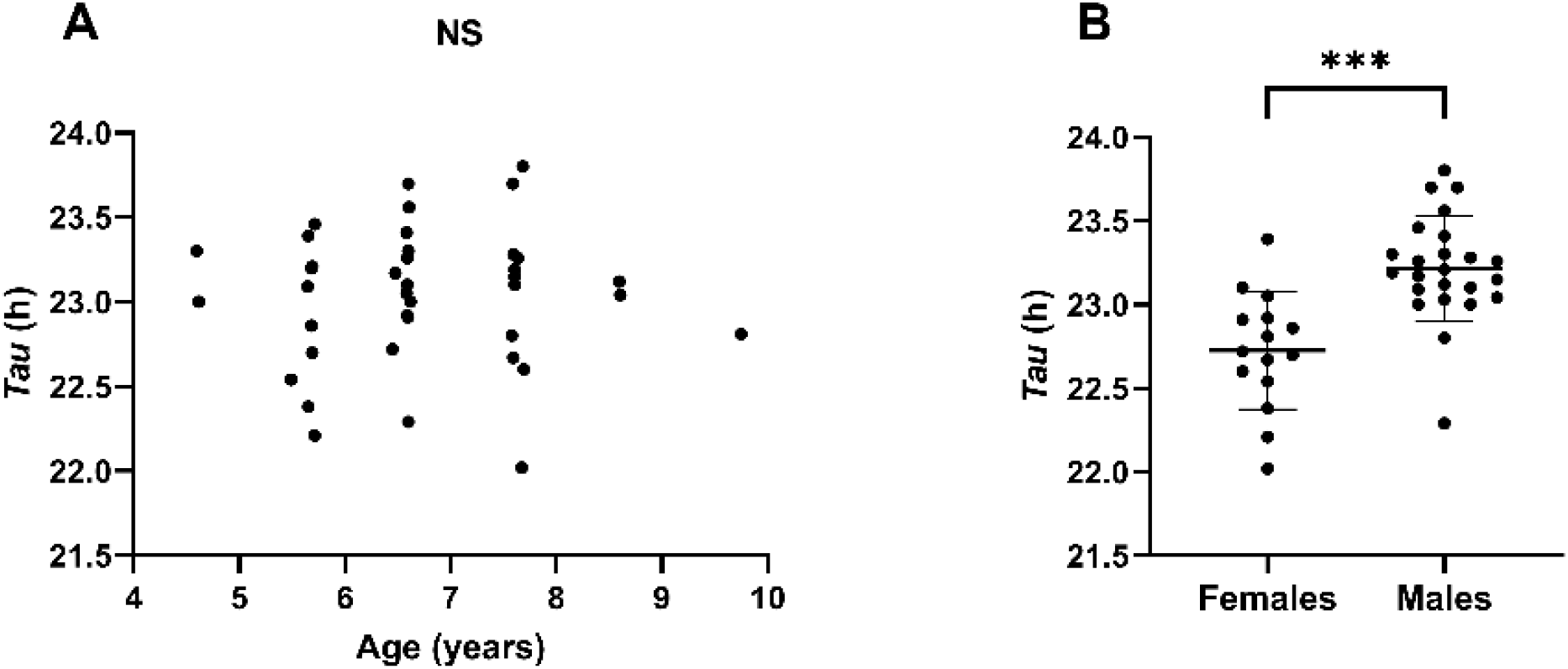
*tau* according to age (A) and sex (B) in the 39 mouse lemurs tested.

**Figure 2:**
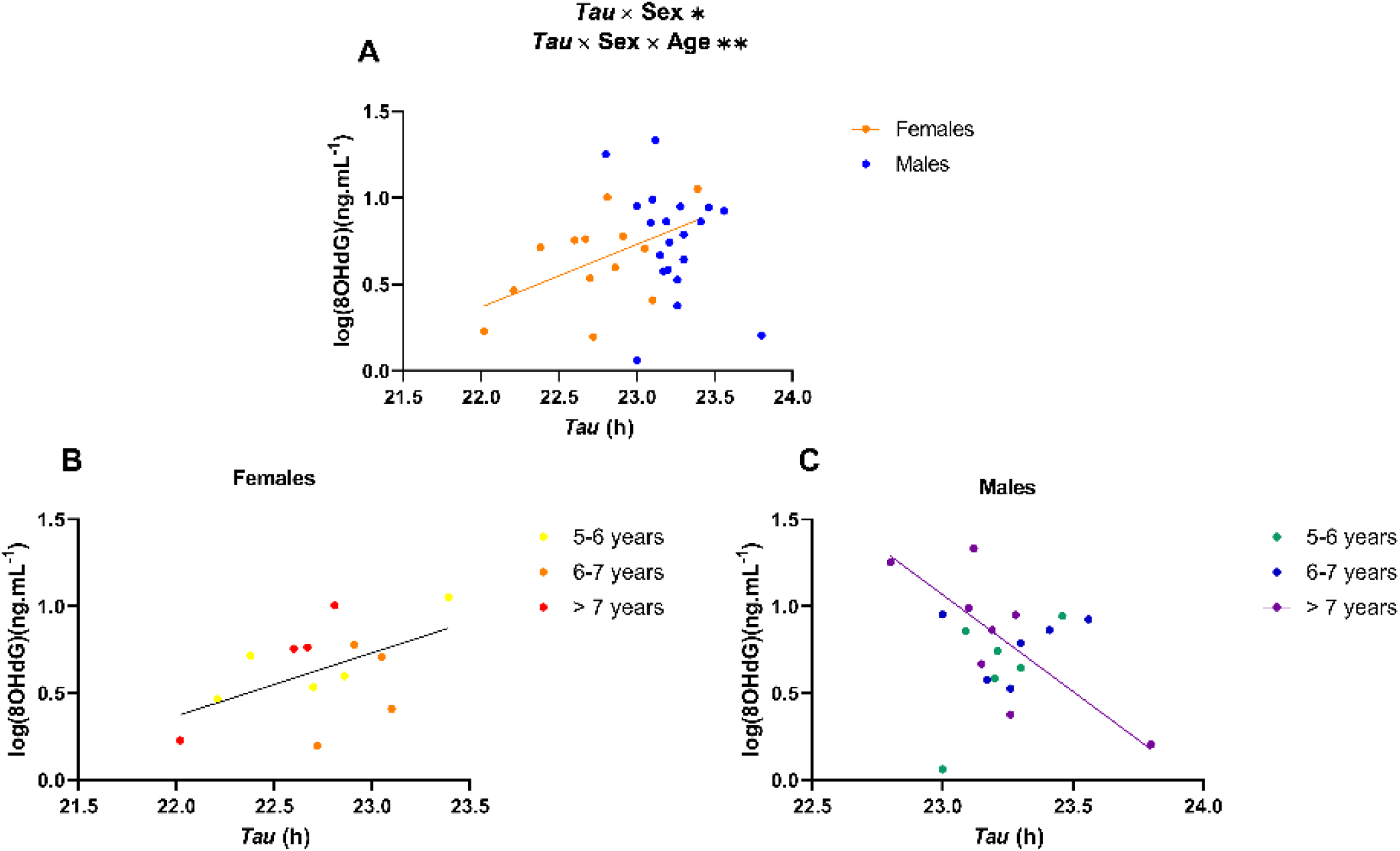
log(8OHdG) plasma levels according to *tau* in females (n=13) and males (n=19) (A) and according to *tau* and age in females (n=13) (B) and males (n=19) (C).

**Figure 3:**
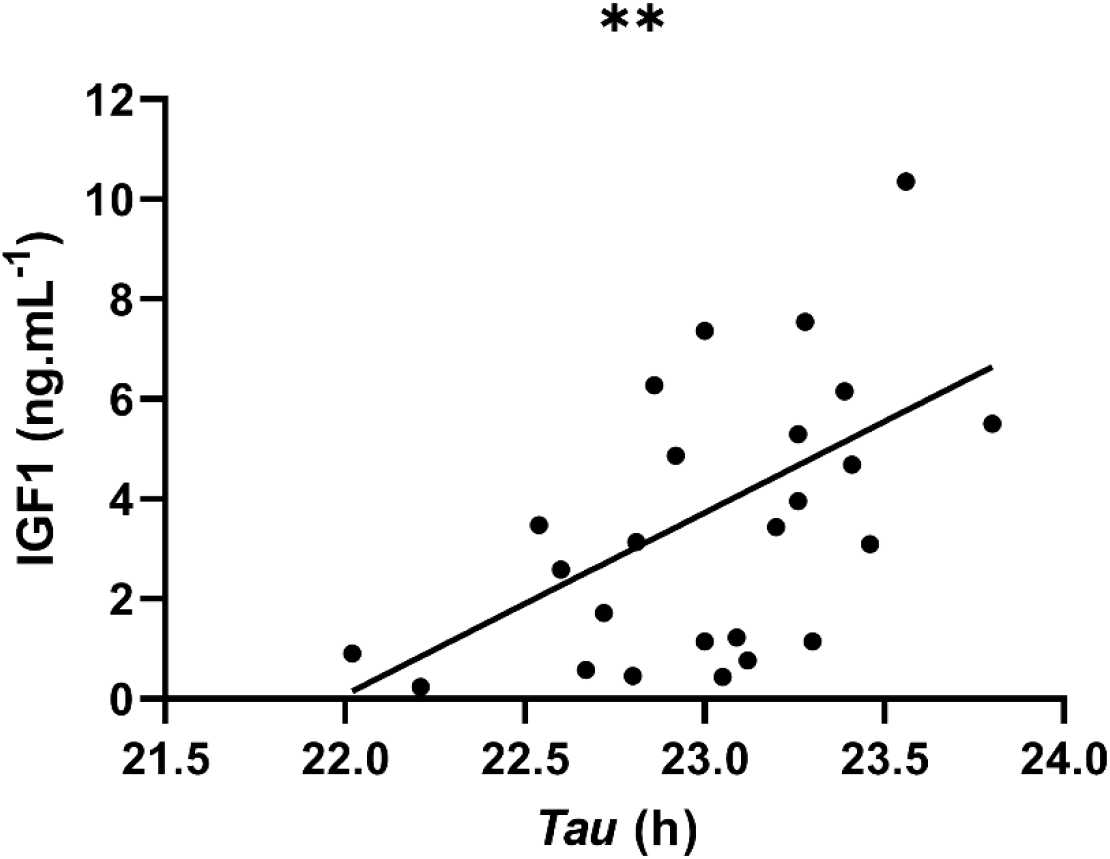
Relationship between IGF-1 plasma concentration and *tau* in 25 individuals.

**Figure 4:**
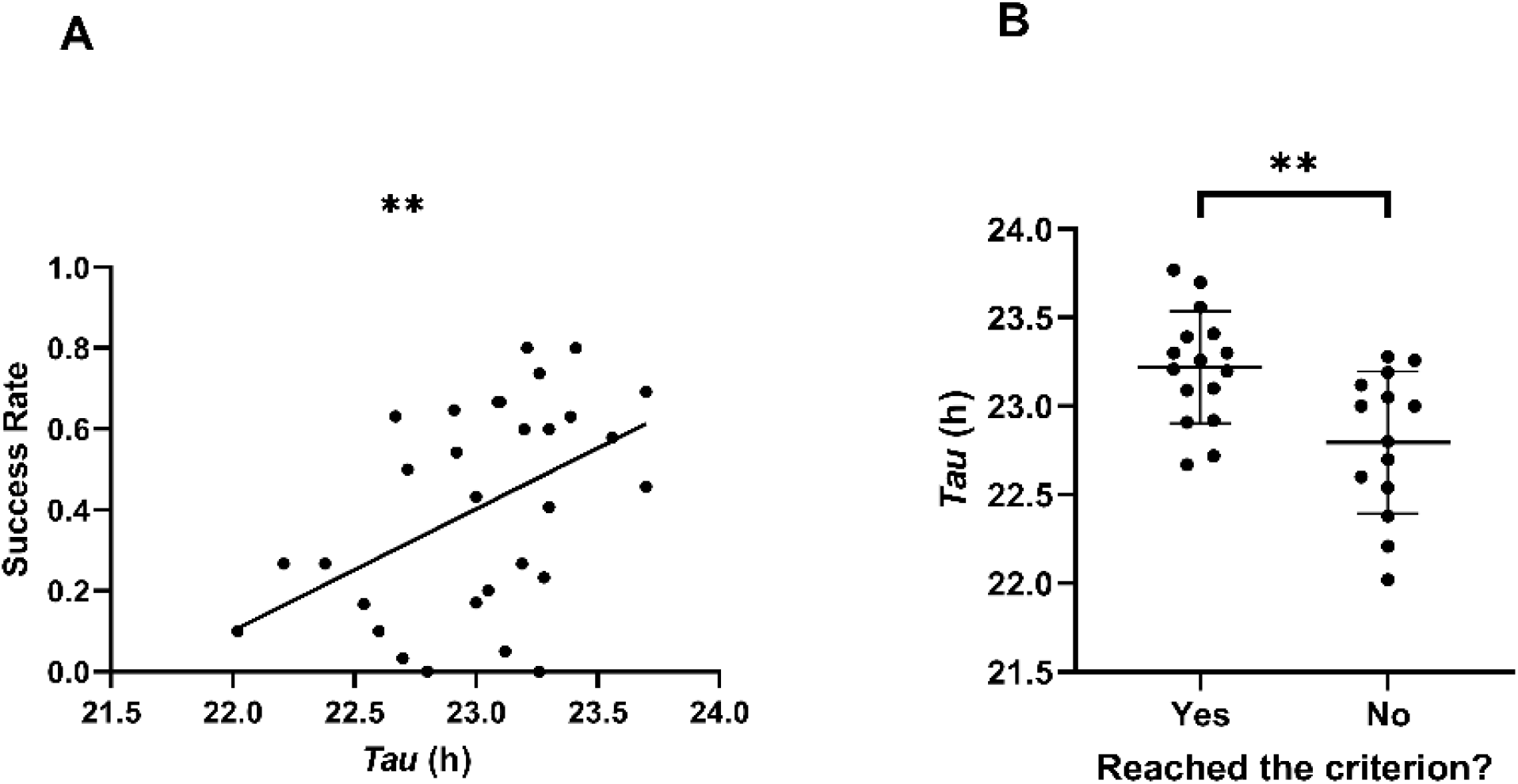
Success rate according to *tau* (n=30) (A) and repartition of *tau* among individuals who reached (n=16) and did not reach (n=14) the success criterion (B).

## Discussion

Our findings revealed that the endogenous period of aged mouse lemurs was significantly correlated with some markers of aging, that might partially explain the underlying mechanisms of the circadian resonance theory. First, *tau* was negatively correlated with 8OHdG concentrations in males over 7 years: in very old individuals, those with an endogenous period closer to 24h were also those with a lower 8OHdG concentration. The oxidative stress level is a validated biomarker of aging, since aging at least partly results from accumulating damages by reactive oxygen species, such as 8OHdG (Niitepõld and Hanski, 2013; Perez-Campo et al., 1998; Russo et al., 2018). Several studies reported an increase in 8OHdG concentrations in humans or rodents after circadian disruption (Grosbellet et al., 2015; Lee et al., 2019), but an effect of *tau* itself had never been reported. The fact that *tau* was only correlated to 8OHdG in male mouse lemurs aged 7 and over may reflect a low impact of *tau* on reactive oxygen species’ accumulation in early aging. However, *tau* may participate in influencing aging in very old individuals by modulating the individual oxidative state. Individuals with *tau* closer to 24h would display lower aging markers and possibly live longer. The positive relationship found in females but not in males is remarkable and contradicts our initial hypothesis. We initially expected that an endogenous period close to 24h would engender less synchronization costs and consequently less oxidative stress in both males and females. This result can related with the results observed in our previous analysis connecting *tau* to survival (Hozer et al., 2020), in which females exhibited no influence of *tau* on survival, whereas male’s survival was strongly correlated to *tau*. The present result could be one of the potential explanations.

In addition, the endogenous period was positively correlated to the IGF-1 plasma concentration without influence of sex. IGF-1 variation in aged population is difficult to interpret because of its pleiotropic effect on biological aging. It is widely accepted that IGF-1 has an important role in aging mechanisms, contributing to age-related pathologies in humans and rodents (Ashpole et al., 2017; Carter et al., 2002; Lind et al., 2019; Mao et al., 2018; Sonntag et al., 1999, 2005). IGF-1 hormone is highly involved in the control of lifespan and aging via changes in specific neuroendocrine pathways, allowing to decelerate growth and preserve resources (Kappeler et al., 2008). On the other hand, IGF-1 has also been noticed for its contradictory actions, more particularly for its beneficial effect in maintaining brain functions during aging. Indeed, direct administration of IGF-1 analogues lead to improve learning and memory tasks and even reverse some effects of cognitive decline (Frater et al., 2017). Bennett and Ingram (2000) found that antagonism of IGF-1 action in the brain impaired both learning and memory. In our study, these observations are corroborated by the fact that the success rate at the cognitive task was positively correlated to the IGF-1 concentration (R=0.43, p=0.04). It has been shown in a previous study in the gray mouse lemur that IGF-1 plasma levels exhibited huge seasonal variations between short-day and long-day seasons, and that short-day plasma concentrations of IGF-1 decreased continuously from the fourth short-day season until death. IGF-1 concentration at the age of four was also positively correlated with survival, making the IGF-1 hormone a good lifespan biomarker in the gray mouse lemur (Aujard et al., 2010). The relationship that we have found between *tau* and IGF-1 concentration suggests that individuals with *tau* far from 24h exhibit a more aged phenotype.

In accordance with the above-mentioned results, we found a positive correlation between learning performances, and *tau*: individuals that reached the success criterion and displayed a higher success rate were those with an endogenous period closer to 24h. In a previous study, we had already found a link between the circadian clock and learning aptitudes: mouse lemurs submitted to a 26h photoperiod exhibited lower performances at the discrimination task than mouse lemurs kept under 24h photoperiod (Hozer and Pifferi, 2020), but to our knowledge, this is the first time that a direct correlation between circadian period and cognitive performances is shown. Since aging in gray mouse lemurs is characterized by moderate to severe decline in executive functions (Languille et al., 2012), the present results are thus consistent with an accelerated aging of cognitive functions when the endogenous period deviates from 24h. However, during normal aging in gray mouse lemur, studies rather report an impairment of memory, while learning is impacted only in very old individuals (Languille et al., 2015; Picq, 2007; Picq et al., 2012). In the context of neurodegenerative aging, though, learning can be altered earlier. For example, mouse lemurs inoculated with Alzheimer brain homogenates displayed premature learning impairments (Gary et al., 2019). This suggests that animals with *tau* very far from 24h exhibit early cognitive decline, as in neurodegenerative aging. The effect of *tau* on learning performances could be attributed to a sleep debt daily generated by the small misalignment between the circadian rhythm and the external periodicity. Indeed, it is well documented that the circadian system mediates cognitive performances such as mood, learning, time-place association or memory in laboratory mice (Albrecht, 2017; Legates et al., 2012; Ruby et al., 2008), including through the direct or indirect implication of sleep (Kyriacou and Hastings, 2010; Sun et al., 2018), a feature that also displays alterations during aging in the gray mouse lemur (Pifferi et al., 2011). Nevertheless, *tau* was not related to spatial working memory in the spontaneous alternation task, whereas several authors reported impaired spatial memory in arrhythmic hamsters or phase-shifted rats (Devan et al., 2001; Ma et al., 2007). However, these latter studies described effects on long-term spatial memory, whereas spatial working memory is a very instantaneous and non-consolidated memory. Only one study showed a decrease in spontaneous alternation in rats submitted to constant light compared to normal photoperiod conditions (Ruby et al., 2013). In mouse lemurs, however, the endogenous period in itself does not seem to influence working memory.

We found no significant relationship between *tau* and metabolic parameters such as dioxygen consumption or energy expenditure, which could seem contradictory with a recent study in which mouse lemurs kept under light-dark cycles of 26h (*i.e.* cycles far from their endogenous period) displayed significant higher VO2 and Heat than control animals kept under 24h photoperiod (Hozer and Pifferi, 2020). Whereas the misalignment between *tau* and external light-dark cycles was at least 2h in that previous study, it was significantly smaller in the present study, since almost all individuals displayed an endogenous period longer than 22.5h. The marginal increase in metabolic rate that we observed in our previous study was maybe undetectable in the present sample.

Interestingly, the endogenous period of females of our sample was significantly shorter than the males’ one. This results echo some previously reported sex differences in endogenous period, with females exhibiting a shorter period than males (humans: (Duffy et al., 2011; Eastman et al., 2017), rodents: (Krizo and Mintz, 2015; Schull et al., 1989). As an explanation for such sexual divergence, authors suggest a neuroendocrine influence, as it has been shown that sexual hormones can modulate several features of circadian responses including the endogenous period (Yan and Silver, 2016). For instance, higher estrogen levels are associated with a shortening of *tau* (Morin et al., 1977). However, this sexual difference had never been reported in the gray mouse lemur so far, even in a recent study comparing several tens of individuals, in which mean endogenous period were 23.10 ± 0.64h in females, which is much higher than in the present study (Hozer et al., 2020). This sexual divergence may be due to a sample effect, but the strong statistical significance does not support this hypothesis. Otherwise, it may lie in the fact that age affects differently males’ and females’ endogenous period. By way of comparison, mean female age was 6.78±1.16 years in the present study *vs* 3.81±2.20 years in the sub-mentioned study (and 6.63±1.08 years and 3.10±1.85 years for males respectively). However, no age effect was found neither in females nor in males in our previous study (Hozer et al., 2020). Further studies will be required to characterize the sex difference in endogenous period and understand its physiological basis, particularly in the gray mouse lemur in which studies are mainly conducted in males (Aujard et al., 2006, 2007; Cayetanot et al., 2005).

In this study, we chose to target metabolic, cellular and cognitive parameters that are known to change with aging in the gray mouse lemur (Aujard et al., 2006, 2010; Hozer et al., 2019; Languille et al., 2012; Marchal et al., 2013; Perret and Aujard, 2006; Picq, 1993; Picq et al., 2015). Most of the measured parameters did not exhibit an age-related change, excepted the cognitive performances that displayed a strong decline with age. From our opinion, this absence of time effect is maybe due to a too small age-window of analysis (most of animals were tested over a range of 2 years, *i.e.* between 6 and 8 years old), leading to relatively weak modifications, that are not representative of the real dynamics of the measured parameters across the whole aging period. However, this hypothesis must be nuanced, since several longitudinal studies in the gray mouse lemur pointed out age-related alterations in immune response, oxidative stress or metabolic rate over two consecutive years (Cayetanot et al., 2009; Marchal et al., 2013; Perret and Aujard, 2006). In conclusion, this study provides interesting insights into the relationships between endogenous circadian period and aging.

By measuring several parameters that are known to vary with aging in the gray mouse lemur, we showed that only some of them correlated with the endogenous period. This suggests that the influence of *tau* on aging processes could only partially explain the mechanisms of the circadian resonance theory. This work calls for further studies on a larger time-scale and more aging biomarkers (such as telomere length), to better assess the link between *tau*, aging and survival.

## Acknowledgments

We are very grateful to Alexxai Kravitz and Bridget Matikainen-Ankney for the development of the infrared sensors system and their availability.

## Authors’ contribution

C.H. and F.P. designed the experimental protocol. C.H. made the experiments and wrote the first draft of the manuscript. C.H. and F.P. reviewed the manuscript.

